# Heterogeneous self-couplings enable scale-invariant memory and cognitive flexibility without criticality

**DOI:** 10.64898/2026.10.25.761657

**Authors:** Francesco Damiani, Gabriele Di Antonio, Jacopo Epifanio, Gianni Valerio Vinci, Andrea Galluzzi, Maurizio Mattia

**Affiliations:** Department of Engineering, Pompeu Fabra University, 08018 Barcelona, Spain; Department of Neuroscience, Istituto Superiore di Sanità, 00161 Rome, Italy

## Abstract

The hierarchical organization of the brain underlies its ability to process information across multiple time scales and support flexible behavior through context-dependent computation. Here, we study a recurrent neural network model with heterogeneous self-interactions across nodes, capturing key features of neuronal circuits, including intrinsic diversity of dynamical time scales and multistable low-dimensional manifolds. We investigate the emergence of scale-invariant dynamics without assuming criticality, challenging the view that power-law statistics necessarily arise from self-organized critical states. We show that heterogeneous self-interactions alone are sufficient to generate scale-invariant memory in subcritical regimes, while enhancing the network’s dynamical richness and computational capacity. Notably, this mechanism promotes the spontaneous emergence of hierarchical organization. We assess the functional implications of this framework using short-term memory tasks and the Wisconsin Card Sorting Test, which probes cognitive flexibility and adaptive rule switching. Compared to models with heterogeneous time constants, our approach achieves comparable performance with more parsimonious synaptic modifications, while preserving scale-invariant dynamics.

## 1 Introduction

The hierarchical and modular organization of the brain is fundamental to its ability to process complex information and adapt to changing environments ^1,2^. Modularity within cortical systems ^3^ and across cortical regions ^4^ emphasizes the importance of structural and functional specialization in cerebral networks. This modularity not only underpins biological neural systems but also informs computational models in artificial intelligence ^5^. In the primate cortex, specialization and hierarchy ^6^ likely stem from diverse neuronal time scales enhancing cortical coordination ^5–8^ and allow the brain to integrate sensory inputs rapidly while generating appropriate context-dependent behavioral responses ^9–15^. These time scales aid complex processing, such as in language comprehension ^8^. Evidence suggests that organized heterogeneity in intra-cluster connectivity^5,6,16-21^ might enable neuronal circuits to exhibit and tune distinct time scales. Understanding how these time scales are modulated and how connectivity patterns are structured to drive behavior remains a significant challenge in neuroscience.

Such structure may reflect an adaptation to environmental complexity, enabling coherent processing and behavior across widely varying and unpredictable inputs. For example, the duration of a spoken sentence can vary substantially with the speaker’s pace, yet its meaning is preserved through hierarchical integration over multiple temporal scales in the cortex ^22–24^. Similar flexibility is required in motor behavior, where actions must be organized into sequences of variable duration and order according to the current goals. Understanding how the brain represents and controls such serial structure remains a central question in motor and cognitive sequencing ^25^. Consistent with this view, preparatory activity in premotor, pre-SMA, and prefrontal cortex reflects sequence order and timing across variable delay periods ^26,27^, while competing actions are selected in a context-dependent manner ^28^.

To process diverse stimuli across changing contexts and adapt behavior accordingly, neural circuits may exploit structural heterogeneity to generate a broad repertoire of internal states and timescales. When the temporal structure of incoming stimuli varies or is not known in advance, relying on a small set of characteristic timescales may be limiting, as circuit dynamics would need to adjust to the temporal statistics of each input. Scale-invariant dynamics provides a possible solution, because it retains information across a broad range of delays without privileging any single characteristic timescale. In such a regime, memory traces decay as a power law, allowing past inputs to influence ongoing dynamics over multiple temporal scales and potentially reducing the need to retune network dynamics to each input ^29^. We hypothesize that structural heterogeneity can generate this form of scale-invariant memory and thereby support flexible computation in environments characterized by variable and unpredictable timescales.

Critical behavior, which is intrinsically associated with scale invariance, has been widely observed in neural systems ^30–33^ and psychophysical studies ^34^, and has motivated the hypothesis that the brain operates near a critical point ^31,32,35,36^.The criticality hypothesis proposes that neural systems function close to the boundary between order and chaos, potentially balancing flexibility and stability and enhancing computational performance ^29,37^. Neural systems could, in principle, self-tune toward such a regime through self-organized criticality ^36,38^. However, this interpretation remains debated, since power-law behaviors can arise in complex dynamical systems without requiring criticality ^39–42^. This raises a fundamental question: can structural heterogeneity generate scale-invariant dynamics and the associated computational flexibility without requiring criticality?

Here, we address this question in heterogeneous recurrent neural networks (RNNs), asking how diversity in local dynamics shapes both the temporal organization of network activity and its capacity for context-dependent computation. Standard RNNs typically assume homogeneous nodes, limiting their ability to capture the dynamical richness of biological networks ^43,44^. Introducing non-negligible self-interactions expands the dynamical repertoire of cortical networks and alleviates catastrophic forgetting in associative memory models ^45,46^, while heterogeneous self-interactions have been shown to broaden the spectrum of spontaneously expressed timescales ^47^. Building on this framework, we show that a specific distribution of self-interaction strengths generates a broad spectrum of timescales and power-law correlations without tuning the network dynamics to be critical. This scale-invariant temporal organization expands the computational range of the model, supporting information processing across multiple timescales. Crucially, the resulting multistability enables rapid adaptation to changing task demands, as illustrated by performance on the Wisconsin Card Sorting Test (WCST), where successful behavior requires maintaining context while flexibly switching rules over variable delays.

## 2 Results

### 2.1 Scale-invariant memory from heterogeneous self-couplings in subcritical RNNs

Building on prior theoretical and empirical work, we ask whether heterogeneous intra-cluster connectivity can generate a wide repertoire of spatiotemporal activity patterns^6,16-18,21,45-47^. To address this question, we introduce a heterogeneous RNN (hRNN) in which heterogeneity arises from unit-specific synaptic self-interactions. The modeled network is composed of *N* units, each representing a homogeneous local population of spiking neurons. Under a mean-field approximation ^48–52^, the activity of each unit is described by a low-dimensional firing-rate dynamics *r*_*i*_ (*t*):

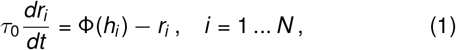

where *τ*_0_ is a uniform relaxation time constant, and Φ*(h*_*i*_*) =* tanh*(h*_*i*_) is the so-called input-output gain function (see Methods). The total input received by unit *i* is

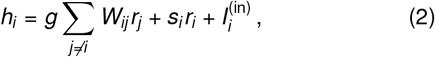

which combines recurrent synaptic input from other units,

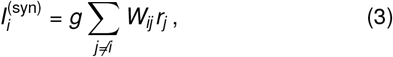

with a unit-specific self-interaction *W*_*ii*_ = *s*_*i*_, and an external input 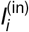 representing signals from other brain regions or environmental stimuli. The synaptic weights *W*_*ij*_ for *i ≠ j* are independently drawn from a zero-mean Gaussian distribution with variance 1*/N* ^53^, while *g* is a global scaling factor determining the strength of inter-unit interactions.

By introducing heterogeneous self-interactions *s*_*i*_ (Fig. 1a), we obtain a network in which different units express markedly different intrinsic timescales (Fig. 1b). Units with sufficiently strong self-coupling, approximately *s*_*i*_ *≳* 1, exhibit bistable dynamics, whereas units with weaker self-coupling, approximately *s*_*i*_ *≲* 1, display monostable or chaotic activity, with the precise transition depending on the global coupling strength *g*^45^,^47^. These regimes are consistent with the coexistence of bistable, chaotic, and monostable dynamics reported in experimental and theoretical studies of cortical activity^16^,^18^,^21^,^54–57^. Importantly, increasing *s*_*i*_ effectively slows the dynamics (Fig. 1b, center; Fig. 1c,d).

**Figure 1.**
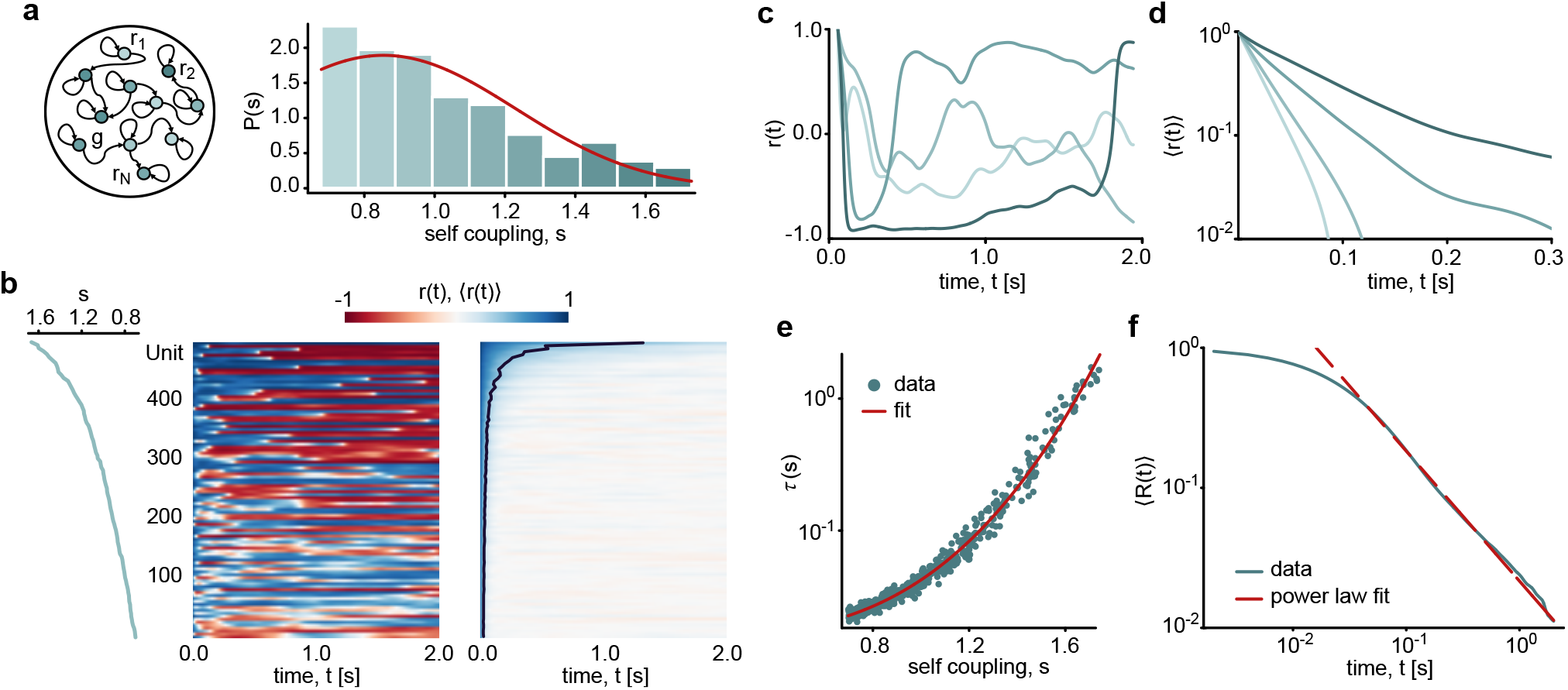
Heterogeneous self-couplings generate multiple timescales and scale-invariant memory in a recurrent neural network. **a**, Schematic of the heterogeneous RNN and distribution of self-couplings *s* used to obtain scale-invariant memory in a network of *N = 500* units. The red curve shows the theoretical distribution *P(s)* derived from the fitted relationship between self-coupling and relaxation time to generate a target power-law decay with *γ = 1* (see Methods). **b**, Left: Self-coupling values *s* assigned to each unit. Center: Raster plot showing the activity of every fifth unit in a single network realization, with all units initialized in the polarized state *r*_*i*_*(0) = +1*. Right: Mean activity ⟨*r*_*i*_(*t*)⟩ of each unit, averaged over 500 realizations of the synaptic matrix with fixed self-couplings and the same polarized initial condition. The black line marks the time at which each unit’s mean activity has decayed by 60% of its initial value. Units with larger self-couplings exhibit longer decay times. **c-d**, Activity traces of representative units with different self-couplings, *s = 0*.*70, 0*.*80, 1*.*00, 1*.*23*, with darker colors indicating larger *s*. **c**, Single-trial activity. **d**, Activity averaged over 500 realizations of the synaptic matrix and plotted on a semi-logarithmic scale to highlight exponential relaxation. **e**, Characteristic decay times *τ* as a function of self-coupling *s*, obtained by fitting the average activity decay of each unit with an exponential function. The red curve shows the fitted relationship 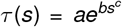 used to derive the distribution *P(s)* in panel a (see Methods). **f**, Population activity ⟨*R*(*t*)⟩, obtained by averaging over all units and 500 realizations of the synaptic matrix. A power-law fit *y = αx*^*− γ*^ is overlaid. To estimate the scaling exponent, the initial transient *t < 0*.*07 s* is excluded, and data beyond *t = 0*.*9 s* are omitted to avoid fitting fluctuations as the activity approaches zero. The coefficient of determination over the fitted interval is *R*^2^ *= 0*.*99*. Fitted parameters are *α = 0*.*0216 (0*.*0001)* and *γ = 0*.*930 (0*.*002)*, with standard errors in parentheses. The fitted exponent is close to the target value *γ = 1*; minor deviations arise from the finite range of available timescales.

When the entire network is initially polarized to *+*1, the mean activity decay ⟨*r*_*i*_*(t)⟩* of each unit – averaged over different realizations of the synaptic weights, with fixed self-couplings – depends on its level of self-coupling, at fixed *g* (Fig. 1b, right). Throughout the paper, we use ⟨·⟩ to denote an average over different realizations of the synaptic matrix. This decay follows an exponential form 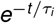 (Fig. 1d), where *τ*_*i*_ is a characteristic timescale (see Methods for further details).

Therefore, adjusting the self-coupling strengths allows us to control the network’s timescales. Figure 1e illustrates the relationship between the decay times *τ* and the values of *s*, demonstrating that increased self-coupling results in slower decorrelation times. Once the relationship *τ(s)* has been established from the extracted single-unit decay times, the distribution of self-couplings *s* can then be adjusted to shape the distribution of timescales *τ* (Fig. 1e,a). Since the mean network activity,

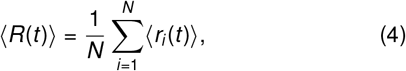

can be expressed as an average of exponential decays with different characteristic timescales, choosing the appropriate distribution of *s* allows us to obtain a target population-level decay (see Methods). Specifically, we identify a parametric distribution *P(s)* for the self-couplings (Fig. 1a) that enables the network to display scale-invariant dynamics, with ⟨*R(t)*⟩ following a power-law decay *βt*^− *γ*^ (Fig. 1). As a proof of concept, we set *γ = 1*, yielding extended memory characterized by a power-law distribution of decay times, *P*_*τ*_*(τ) ∝ τ*^−2^, consistent with experimental observations^58^. This approach can be generalized to other values of *γ*, including *γ = 3/2*, which is commonly observed in critical phenomena^59^.

To characterize the asymptotic dynamical regime underlying this scale-invariant relaxation, we performed long simulations across independent network realizations, starting from the polarized initial condition (*r*_*i*_*(0) = +1* for all units) and lasting up to 10^4^ s (see Methods). The vast majority of these realizations(93%) converged to stable fixed points within the simulated interval, following prolonged transients (Fig. S1a,b), while the remaining 7% exhibited dynamics consistent with stable periodic attractors. For the fixed-point realizations, the largest Lyapunov exponent was negative (Fig. S1c), indicating asymptotically stable, subcritical dynamics. We further assessed the robustness of this regime by varying the recurrent gain *g* by ±5% around its reference value (*g* = 0.4). Across this range, the largest Lyapunov exponents remained non-positive for almost all network realizations, providing no evidence for a qualitative change in the asymptotic dynamical regime (Fig. S1c).

Together, these results show that local recurrent heterogeneity can be appropriately shaped to combine multiple dynamical regimes and intrinsic timescales into scale-invariant memory of past stimuli at the population level, without requiring criticality.

### 2.2 Noise-driven origins of a broad timescale repertoire

To clarify the link between local intra-cluster connectivity and the intrinsic timescales expressed by the network, we provide here a mechanistic interpretation of the *τ(s)* relationship. We show that, for units in the bistable regime^45^ (those with *s*_*i*_ *> 1*), the emergent timescales can be effectively described as noise-driven escape times of a Brownian particle in a double-well potential, where thermal fluctuations are equivalent to a colored noise^61^.

In the hRNN off-diagonal recurrent interactions generate fluctuating input currents that perturb units with different self-couplings. These currents are not expected to be independent Ornstein-Uhlenbeck (OU) processes, since their correlation structure arises from a recurrent system. Nevertheless, if their main role is to provide an effective stochastic drive, then a fully decoupled network with the same heterogeneous self-couplings should approximately recover the same memory properties when driven by colored noise of appropriate strength and correlation time. To test this hypothesis, we compare the original coupled network with a decoupled network in which all off-diagonal recurrent interactions are removed, while preserving the same self-couplings (Fig. 2a). In the coupled network, each unit receives the recurrent input

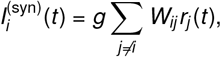

whereas in the decoupled network the recurrent input to each unit is replaced directly by an independent Ornstein– Uhlenbeck (OU) process,

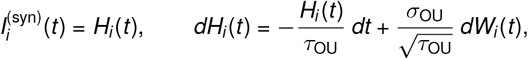

where *H*_*i*_*(t)* is the OU process driving unit *i, τ*_*OU*_ and *σ*_*OU*_ determine its correlation time and amplitude, respectively, and *dW*_*i*_*(t)* are independent Wiener increments.

**Figure 2.**
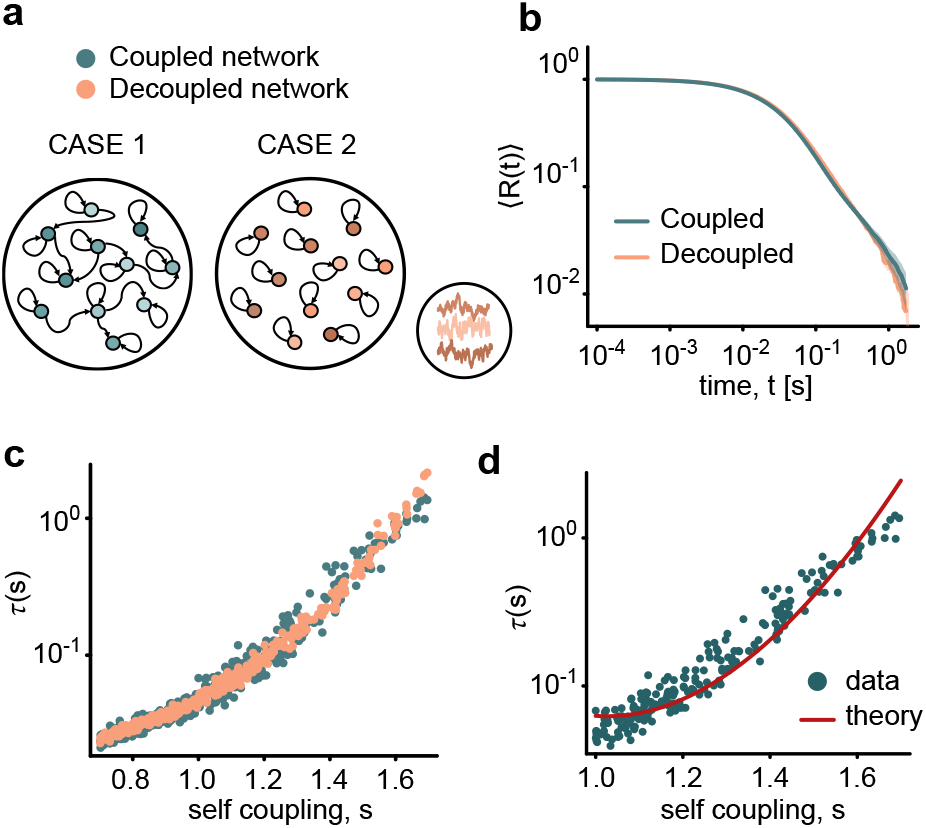
Effective colored noise reproduces recurrent synaptic input and scale-invariant memory. **a**, Schematic of the two network configurations. Case 1: the coupled heterogeneous RNN introduced in Fig. 1. Case 2: a decoupled heterogeneous network with the same self-couplings {*s*_*i*_*}*, in which off-diagonal recurrent interactions are removed and synaptic currents are replaced by independent Ornstein-Uhlenbeck (OU) processes. **b**, Mean population activity ⟨*R(t)*⟩ after initialization from a polarized state (*r*_*i*_*(0) = +*1 for all *i*), shown for the coupled network and the OU-driven decoupled network. **c**, Characteristic times *τ(s)* extracted for individual units in the coupled and decoupled networks. **d**, For bistable units, *s > 1*, relaxation times extracted from the coupled network are compared with the theoretical escape-time prediction for bistable rate units driven by colored synaptic noise^60^ (see Methods).

This reduction is motivated by dynamic mean-field descriptions of random recurrent networks^45^,^53^. An exact dynamic mean-field treatment of the present heterogeneous network would require a nontrivial self-consistent description of recurrent input statistics across units with different self-couplings and broad temporal correlations. We therefore adopt a minimal effective-noise approximation, in which the recurrent input is replaced by an externally generated colored stochastic drive with tunable amplitude *σ*_*OU*_ and correlation time *τ*_*OU*_. The OU process should therefore not be interpreted as an exact description of the recurrent currents, but as a reduced description that tests whether their leading functional role is to provide fluctuations of the appropriate magnitude and temporal structure.

We set the OU correlation time equal to the single-unit time constant, *τ*_*OU*_ *= τ*_0_ *= 0*.*01 s*, matching the condition under which analytical predictions for noise-driven escape times in bistable neuronal populations are available^60^. With *τ*_*OU*_ fixed, the noise amplitude *σ*_*OU*_ is the only remaining free parameter. We fit it to reproduce the population-level relaxation of the coupled network from the same polarized initial condition (see Methods).

Despite the complete removal of recurrent interactions, the OU-driven decoupled network closely reproduces the scale-invariant relaxation of the coupled network (Fig. 2b). The agreement also holds at the single-unit level: the characteristic relaxation times *τ(s)* extracted from the decoupled network remain close to those extracted from the coupled network across the range of self-couplings (Fig. 2c).

This indicates that the hierarchy of timescales is primarily shaped by the heterogeneous self-coupling values, while recurrent interactions act, to leading order, as a source of fluctuations that drive transitions between activity states. This interpretation is especially clear in the bistable regime, *s > 1*. In this range, each unit can be approximated as a bistable dynamical system driven by colored noise: the self-coupling *s* controls the stability of the two activity states, whereas the amplitude of the input fluctuations controls the probability of noise-driven transitions. Consistently, the relaxation times extracted from the coupled network are well captured by the theoretical prediction for escape times in bistable rate units driven by colored synaptic noise^60^ (Fig. 2d; see Methods).

As an independent consistency check, the fitted OU amplitude yields fluctuations of comparable magnitude to the recurrent currents in the original network, 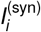. Specifically, *σ*_*OU*_ *≃ 0*.*41* corresponds to a stationary current standard deviation 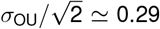, close to the value *≃ 0*.*27* measured in the coupled network (see Methods).

Together, these results support an effective description in which heterogeneous self-couplings set a broad distribution of local relaxation timescales, while recurrent interactions primarily provide stochastic fluctuations that drive the relaxation dynamics. Although the recurrent currents are neither independent OU processes nor characterized by a single timescale, the effective-noise description is sufficient to reproduce both the heterogeneous local timescales and the resulting scale-invariant population decay. Notably, this noise-driven mechanism can, in principle, support arbitrarily long memory timescales simply by shaping the distribution of self-couplings *s*_*i*_, without requiring correspondingly slow timescales in the microscopic dynamics of individual units.

### 2.3 Scale-free short-term memory

Having established that heterogeneous self-couplings can generate a broad repertoire of intrinsic timescales, we next ask whether this temporal organization confers a functional advantage. Stimuli from the environment arrive at variable, often unpredictable rates: a circuit tied to a single characteristic timescale would need retuning to match each input rate, whereas a scale-free memory should handle a broad range of inter-stimulus delays *T*_sep_ with fixed dynamics. We therefore test whether our hRNN can retain structured inputs robustly across different temporal intervals, leveraging the coexistence of multiple intrinsic timescales within a single network without requiring operation near a critical state.

Figure 3a shows the activity of a single hRNN realization over one stimulation cycle, during which six orthogonal stimuli are presented sequentially and labeled from 1 to 6 (see Methods). Each stimulus evokes a peak in the population-averaged activity *R*(*t*) (middle) and a corresponding vertical pattern in the activity raster (bottom), reflecting the simultaneous polarization of the stimulated units. Between successive stimuli, individual units continue to fluctuate across their available activity states, while the population activity relaxes on average toward zero.

**Figure 3.**
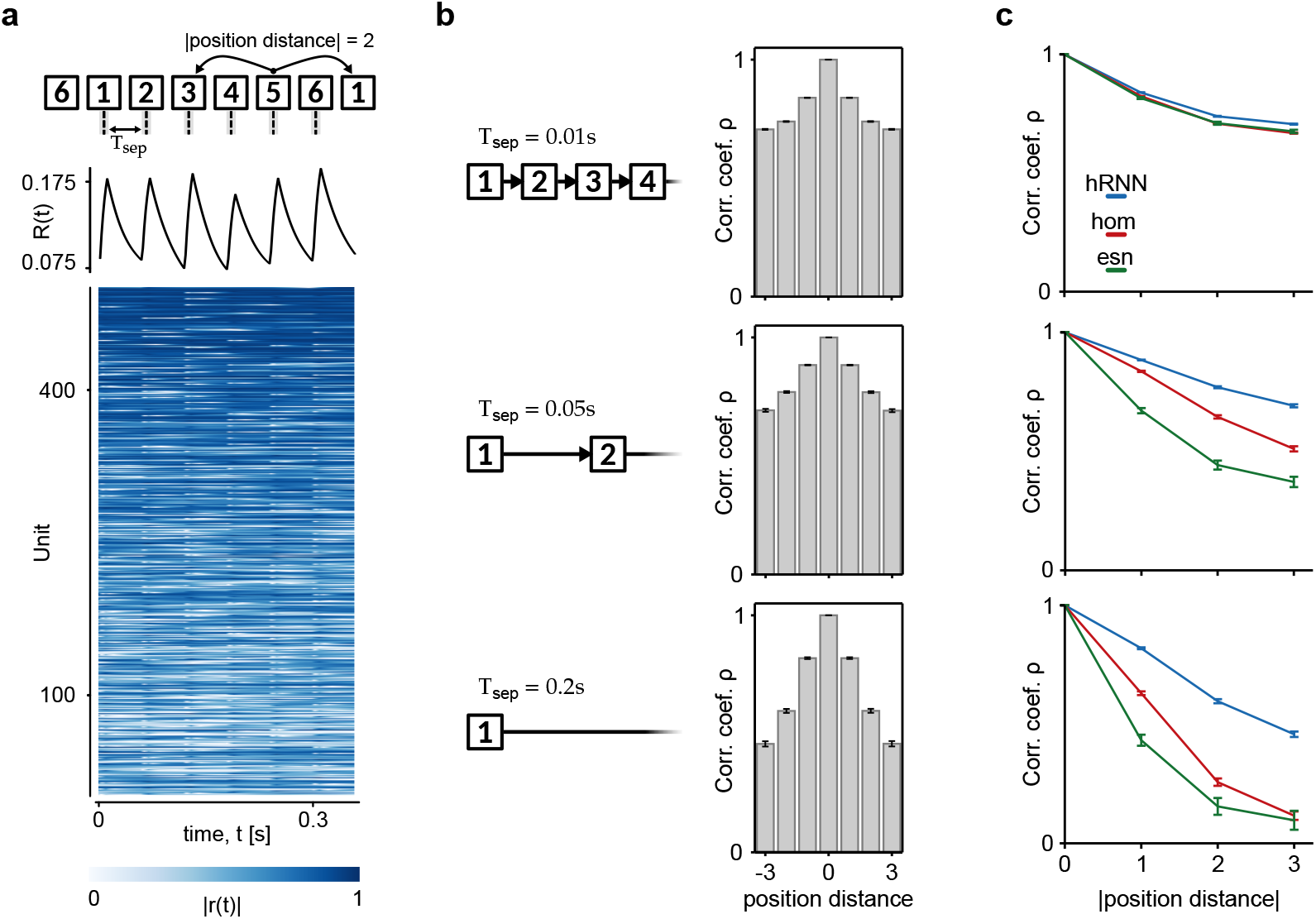
Self-coupling heterogeneity yields delay-invariant short-term memory. **a**, Top: six orthogonal stimuli, evenly spaced in time, each driving a different subset of approximately one sixth of the units toward +1 for 10 ms are presented to the heterogeneous RNN (hRNN). Middle: population-averaged activity *R(t)* of the network. Bottom: raster plot of the absolute unit activities |*r*_*j*_*(t)*|. A single network realization over one stimulation cycle is shown. **b**, Mean Pearson correlation between evoked activity vector in the hRNN as a function of the signed position distance between stimuli (their ordinal separation in the sequence). Each row corresponds to a different inter-stimulus delay *T*_*sep*_ *= {0*.*01, 0*.*05, 0*.*2}* s. **c**, The same correlation, shown for non-negative position distances, compared across the hRNN, the homogeneous network (hom) and the echo-state network (ESN) for the same task as in (a). Results in **b–c** are averaged over 50 synaptic-matrix realizations and 10 consecutive stimulation cycles. Error bars indicate the SEM across realizations.

In each row of Fig. 3b we report the mean Pearson correlation coefficients between the activity vectors evoked by each stimulus, computed as a function of the position distance *d* between stimuli (their ordinal separation in the presentation sequence), averaged over network realizations; each row corresponds to a different value of *T*_sep_. We then compare these results with two alternative network architectures in the same short-term memory paradigm. The first is a homogeneous RNN, identical to the hRNN except that all units share a single self-coupling value, set to the mean of the hRNN distribution. As a result, this network lacks the broad distribution of intrinsic timescales generated by heterogeneous self-couplings. The second is a classical echo-state network (ESN) ^44,62^, with random recurrent connectivity and no self-interactions. Its recurrent gain was set to *g*_ESN_ = 0.9999, at the edge of stability where echo-state networks are classically expected to maximize memory ^37^; the ESN therefore provides a near-critical reference against which the hRNN can be compared. The ESN time constant was chosen to yield correlations comparable to those of the homogeneous network at the shortest inter-stimulus delay, allowing us to assess how the different architectures respond as the input timescale is increased while keeping network parameters fixed (see Methods). As expected, the persistence of past stimulus patterns becomes increasingly sensitive to the stimulation interval. In both the ESN and the homogeneous RNN, correlations decay increasingly fast with position distance as *T*_sep_ grows, approaching zero at distance 3 for *T*_sep_ = 0.2 s. In contrast, the hRNN is substantially more robust to changes in input timing, with correlations decaying more gradually with position distance even at the longest inter-stimulus delay (Fig. 3c). This delay, *T*_sep_ = 0.2 s, is twenty times the hRNN single-unit time constant *τ*_0_ = 0.01 s.

This greater robustness of the hRNN, relative to the homogeneous RNN and the ESN, reflects its broad repertoire of intrinsic timescales. Notably, the ESN operates at the edge of stability, so that long-lasting memory across delays is achieved by the hRNN but not by a near-critical reservoir. Unlike the two homogeneous architectures, the hRNN retains stimulus information across a wide range of stimulation rates. Thus, the co-existence of multiple timescales within a single network makes short-term memory substantially less sensitive to input timing. Heterogeneous self-couplings provide a simple mechanism for incorporating long timescales while, through local bistability, simultaneously enriching the network’s dynamical landscape. We next show that this richer landscape provides a repertoire of internal states that supports context-dependent computation while preserving scale-free short-term memory.

### 2.4 Context-dependent computation in hRNNs

Context-dependent behavior requires the integration of information across multiple timescales, combining rapid processing of current stimuli with the maintenance and updating of latent contextual states. The WCST provides a paradigmatic example of this challenge. In the task version considered here, a subject is presented with a central card and two choice cards that differ in features such as color and shape. The subject must select the card matching either the color or the shape of the central card, according to the currently valid rule. At unpredictable times, the rule changes, requiring the subject to infer the new rule from error feedback and adapt behavior accordingly.

To test whether scale-free memory and multistability support such adaptive behavior, we implemented a simplified WCST within the hRNN. Four cards were defined by two possible shapes and two possible colors. Each card was represented as a square-wave sensory input delivered to the hRNN, with distinct cards projecting onto the network through independent random input weights (Fig. 4a). At the beginning of each trial, presentation of the central card generated a positive input current. After a delay period, two choice cards were presented simultaneously, one associated with a positive input and the other with a negative input representing their position. The network responded by selecting either the right or left option, encoded as a positive or negative output pulse, respectively. The active matching rule (shape or color) changed between trials with a probability of 20%. Following an incorrect choice, an additional error signal was delivered to the network. Under these conditions, the external input to node *i* is given by

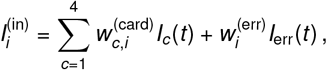

where 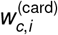 and 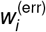 are independently drawn from Gaussian distributions with zero mean and standard deviations *g*_card_ and *g*_error_, respectively.

**Figure 4.**
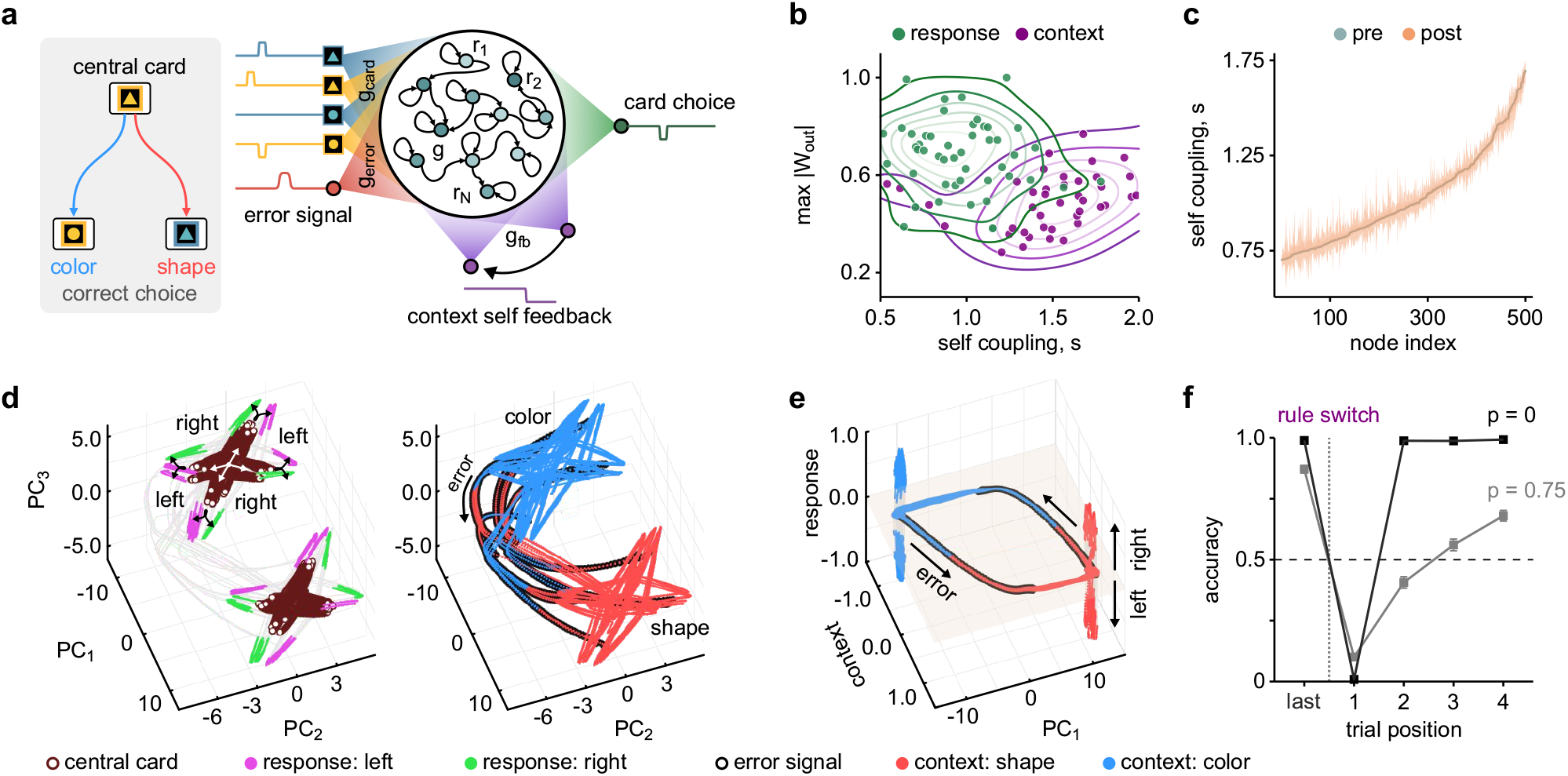
Heterogeneous network strategy for solving the Wisconsin Card Sorting Test. **a**, Schematic representation of a heterogeneous reservoir addressing the WCST, with an inset illustrating the task solution. Given a central card and a pair of oriented cards, the system must respond based on the current context, which changes over time with a certain probability. **b**, The reservoir exploits short timescales for card selection and long timescales for context encoding. Points indicate the maximum readout weights associated with the response (dark green) and context (dark purple) for 50 independently trained reservoirs; overlaid curves show kernel density estimates. **c**, Sorted pre-learning self-coupling values, with a ribbon indicating post-learning variation averaged over 50 networks. **d**, The network selects an attractor for each context (blue, red; right panel). The error signal facilitates transitions from one context to another. Within each attractor, every combination of cards is represented by a fixed point. Initially, the central card shifts the network from its resting fixed point (white arrows, left panel), producing a star-like pattern corresponding to the four possible central cards, before the pair of cards that follows directs the neural state towards either a right response zone (green) or a left response zone (fuchsia). **e**, Hysteresis cycle in the context–PC1 plane during context transition. The response activity develops in a space orthogonal to this plane. **f**, Fraction of correct responses on the last trial before a rule switch and on the first four trials after it. During testing each error signal is halved with probability *p* = 0 (black) and *p* = 0.75 (gray). Vertical dotted line: rule switch; dashed line: chance level. Mean ± SEM over 200 networks, counting only the switches between two blocks of at least four trials. Network parameters as in Fig. 1, with external input intensities *g*_error_ = 1.75, *g*_card_ = 1.0, and *g*_fb_ = 0.8.

The network was trained over 80 trials using the reservoir computing framework for echo-state networks ^62^ (see Methods).Accordingly, learning was restricted to the readout 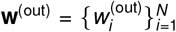, which were fitted by linear regression to map the network state onto the behavioral output, corresponding to the selected card. During training, the hRNN additionally received external context signal 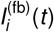, encoded as +1 and − 1 for the shape and color rules, respectively. This signal projected onto the network through random feedback weights 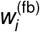 with unit variance modulated by the feedback strength *g*_fb_. In parallel, a second set of linear readout weights, **w**^(ctx)^, was trained to reconstruct the contextual signal from the network activity, such that

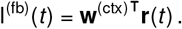

During testing, the external context signal was removed and replaced by its reconstruction from the network state (Fig. 4a), thereby closing the feedback loop and enabling the network to autonomously maintain the latent contextual variable. This procedure is mathematically equivalent to modifying the recurrent connectivity through the rank-one perturbation

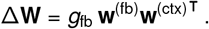

During the test phase, the hRNN successfully solved the WCST while autonomously maintaining the task context through the closed feedback loop. To examine how information is distributed across the heterogeneous population, we analyzed the readout weights associated with card selection and context representation. As shown in Fig. 4b, card-selection readouts are predominantly concentrated on units with small self-couplings, whereas context-related readouts extend toward larger self-coupling values. These results indicate a functional segregation across timescales, with faster units contributing primarily to stimulus-driven decisions, whereas slower and locally multistable units (*s*_*i*_ *>* 1) provide a reservoir of stable states capable of sustaining contextual information. This organization is preserved during autonomous operation, as context feedback induces only minor perturbations of the effective self-couplings (Fig. 4c). The coexistence of fast and slow dynamical modes therefore enables the simultaneous processing of sensory stimuli and maintenance of latent task variables. We return to this mechanism in the next section, where we compare self-coupling heterogeneity with heterogeneity in intrinsic time constants.

The dynamical organization underlying task performance becomes clearer when examining the network state space (Fig. 4d,e). Projection onto the first three principal components of r(*t*) (PC_1_, PC_2_, PC3) reveals two well-separated attractor regions corresponding to the two task contexts (blue and red in Fig. 4d-right). Presentation of the central card displaces the network state away from its equilibrium manifold, after which the trajectory evolves toward either the right-choice (green) or left-choice (fuchsia) response region depending on the presented card pair (Fig. 4d-left). Error feedback induces transitions between contextual attractors, allowing the network to switch behavioral rules. These transitions form a characteristic hysteresis loop in the 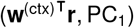 plane (Fig. 4e), illustrating how context-dependent behavior emerges from the multistable structure of the network dynamics.

Interestingly, despite the absence of any explicit constraint promoting abstraction during learning, the resulting geometrical organization of neural activity closely resembles that recently reported in the human hippocampus during inferential reasoning tasks ^63^. In particular, stimulus- and response-related activity occupies dimensions largely orthogonal to those encoding task context, which can be decoded primarily along the PC_1_ axis (Fig. 4d-right). Such a separation of contextual and stimulus-dependent representations has been proposed as a key ingredient for abstraction and generalization ^63^.

Motivated by this similarity in neural geometry, we asked whether the model could also reproduce the variability observed in human behavioral performance. To this end, we introduced variability in the efficacy of the error signal, mimicking differences in the salience or surprise associated with rule violations. Specifically, each error signal was reduced by a factor of two with probability *p*, independently across rule switches (see Methods). When all error signals were delivered at full strength (*p* = 0), the network closely reproduced the behavior of human subjects displaying strong inferential abilities (Fig. 4f, black). Increasing the probability of weakened error signals progressively delayed contextual updating, leading to a more gradual inference of the active rule, similar to the behavior observed in some human subjects ^63^ and non-human primates ^64^.

These results demonstrate that heterogeneous self-couplings naturally support a division of labor across timescales, with fast dynamics mediating stimulus-driven decisions and slower dynamics maintaining contextual information. Importantly, the same heterogeneity can generate a rich repertoire of stable and metastable states associated with units operating in the locally multistable regime (*s*_*i*_ *>* 1). This multiscale and multistable organization gives rise to contextual attractors and enables adaptive behavior through relatively small modifications of the recurrent connectivity.

### 2.5 Beyond timescale heterogeneity: intrinsic multistability

The results above raise a natural question: are the contextual attractors observed in the hRNN simply a consequence of the broad spectrum of timescales generated by heterogeneity, or do they rely on the intrinsic multistable landscape produced by heterogeneous self-couplings? To address this question, we compared the hRNN of Fig. 4 with networks of identical architecture in which self-couplings were homogeneous and heterogeneity was introduced instead through uniformly distributed intrinsic time constants (*τ*_0_-networks; see Methods).

Both variants solve the WCST with the same low-dimensional strategy illustrated in Fig. 4, yet they exhibit markedly different structural changes following learning. Whereas the distribution of self-couplings remains largely unchanged in the hRNN (Fig. 5a), learning induces a substantial departure from the initial delta-function distribution in the *τ*_0_-network, giving rise to an approximately Gaussian distribution of effective self-couplings (Fig. 5b). In both models, performance improves with increasing feedback strength *g*_fb_ over a comparable range of values, following similar trends (Fig. 5c). The similarity of these performance curves indicates that the two architectures were compared under conditions yielding comparable task proficiency.

**Figure 5.**
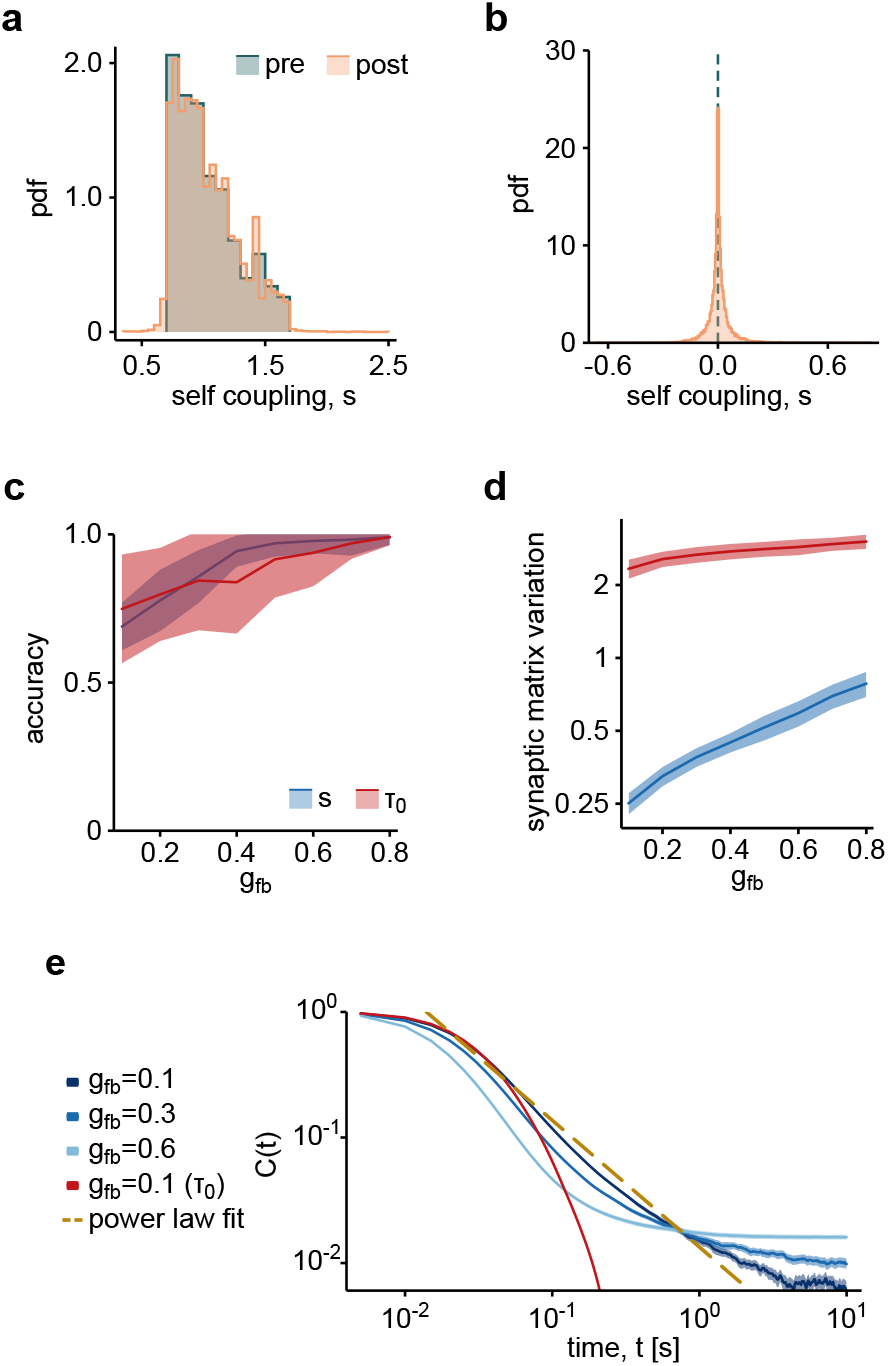
Comparison of sources of heterogeneity. **a–b**, Distributions of self-couplings before (blue) and after (orange) learning in networks with heterogeneous self-couplings *s* (**a**) or heterogeneous time constants *τ*_0_ (**b**). **c**, Change in the fraction of correct responses, excluding rule-switch trials, as a function of context-feedback strength for networks with heterogeneous *s* (blue) or *τ*_0_ (red). **d**, Relative change in the synaptic weight matrix after learning, quantified by the ratio of Frobenius norms ‖ Δ **W** ‖ _F_*/* ‖**W** ‖_F_, as a function of context-feedback strength, shown on a logarithmic scale. Shaded bands indicate the standard deviation across runs. **e**, Autocorrelation functions (mean ± SEM) for different context-feedback strengths. The dashed line shows a least-squares power-law fit, *C*(*t*) = α*t*^*−γ*^, to the *g*_fb_ = 0.1 curve, excluding the initial plateau (*t <* 0.02 s), yielding *γ* = 1.00 ± 0.14. Results are averaged over 50 random networks.

Increased feedback strength is associated with larger modifications of the recurrent connectivity. We quantify these changes through the ratio between the norm of the rank-one (post-learning) correction and that of the original connectivity matrix, ‖ Δ**W** ‖*/* ‖ **W** ‖. Across all tested values of *g*_fb_, the required correction is nearly an order of magnitude smaller in the hRNN than in the *τ*_0_-network (Fig. 5d). This difference suggests that heterogeneous self-couplings provide a pre-existing dynamical scaffold that can be recruited to encode task context. In particular, units operating in the locally bistable regime (*s*_*i*_ *>* 1) endow the network with an intrinsic repertoire of stable and metastable states, whereas in the *τ*_0_-network the contextual attractors must be constructed through substantially larger modifications of the recurrent connectivity.

Although increasing *g*_fb_ can improve task performance, excessive feedback progressively suppresses the spontaneous fluctuations required to preserve scale-free short-term memory, ultimately stabilizing a single contextual state and reducing the probability of spontaneous rule transitions (Fig. 5e). Interestingly, perfect performance is not necessarily desirable in the present setting. Humans and non-human primates frequently commit errors following rule switches because contextual changes must be inferred from feedback ^63,64^, as faithfully replicated by our model (Fig. 4f). Consistent with this observation, the hRNN operates effectively in a regime of weak feedback where power-law memory, spontaneous fluctuations, and adaptive context switching coexist, without requiring a homeostatic mechanism such as self-organized criticality ^36,38^.

These results identify intrinsic multistability, rather than timescale heterogeneity alone, as the key ingredient enabling the coexistence of scale-invariant memory and context-dependent computation. Heterogeneous self-couplings provide a robust alternative to criticality, generating both the temporal richness and the attractor structure required for adaptive cognitive processing.

## 3 Discussion

A longstanding challenge in theoretical neuroscience is to identify network mechanisms capable of simultaneously supporting long memory timescales, flexible computation, and adaptation to changing contexts. In this work, we show that structural heterogeneity in recurrent networks provides a unified solution to these demands. Heterogeneous self-couplings generate both a broad hierarchy of intrinsic timescales and a rich repertoire of stable and metastable states, allowing scale-free memory and context-dependent computation to coexist within the same network architecture. The resulting dynamics combine temporal richness with cognitive flexibility, suggesting that heterogeneity may constitute a general organizational principle for neural computation.

A central question motivating this work was whether scale-invariant dynamics and the associated computational flexibility necessarily require neural circuits to operate near a critical point. The criticality hypothesis proposes that neural systems may achieve optimal computational performance by functioning close to a transition between order and chaos ^29,37^, a view motivated by numerous observations of critical-like phenomena in neural activity^31–33,65,66^. Our results suggest an alternative interpretation. Rather than emerging from proximity to a global critical transition, scale invariance arises from the distribution of local dynamical properties induced by self-coupling heterogeneity. A broad distribution of self-couplings generates a continuum of relaxation timescales and power-law temporal correlations, while simultaneously establishing a multistable landscape capable of supporting context representations. Importantly, the networks considered here remain globally subcritical: despite potentially long transient dynamics, activity ultimately converges to stable attractors. This observation is consistent with previous studies showing that power-law behavior may emerge in complex systems without criticality ^39–42^. Furthermore, learning does not disrupt these scale-invariant properties. In particular, weak contextual feedback preserves both power-law memory and flexible context switching, allowing scale-free dynamics and adaptive computation to coexist. While these findings do not exclude a role for criticality in brain function, they suggest that structural heterogeneity may provide a complementary and potentially more robust route to the emergence of temporal complexity and cognitive flexibility.

Our results suggest a broader interpretation of structural heterogeneity in cortical organization. Rather than viewing individual cortical areas as isolated heterogeneous networks, the cortex itself may be regarded as a hierarchy of heterogeneous recurrent networks differing in the extent of their self-coupling heterogeneity. In this view, the maximum self-coupling strength *s*_max_ = max_*i*_ *s*_*i*_ acts as a coarse measure of the dynamical complexity available to a local circuit. Cortical regions closer to sensory inputs would be characterized by relatively small values of *s*_max_, yielding faster dynamics, weaker multistability, and neural representations closely constrained by incoming stimuli. Progressively higher-order associative areas would instead exhibit larger values of *s*_max_, extending the spectrum of timescales and enriching the repertoire of stable and metastable states available for internal computation. Such circuits would be increasingly capable of integrating information over longer temporal windows and of representing contextual variables that cannot be inferred from instantaneous sensory input alone.

This perspective offers a potential unifying explanation for two prominent organizational principles of the cortex. First, increasing *s*_max_ naturally predicts a rostro-caudal gradient of intrinsic timescales, consistent with proposals that higher cortical areas integrate information over progressively longer temporal windows ^8, 22, 56, 58^. Second, the concomitant increase in multistability predicts a richer and more separable representational geometry, providing a possible dynamical basis for the recently reported increase in representational complexity along the cortical hierarchy ^67^. Within this framework, temporal integration and representational complexity emerge as complementary consequences of the same underlying increase in network heterogeneity. Testing this hypothesis would require estimating local dynamical heterogeneity across cortical areas and assessing whether it covaries with both intrinsic timescales and representational dimensionality.

The present results also suggest a possible dynamical mechanism underlying behavioral flexibility and, more generally, learning-to-learn. In our implementation of the WCST, successful performance emerges through the organization of neural activity around a set of contextual attractors that are accessed through context-dependent responses to external events, such as error feedback. Notably, this solution recovers key aspects of previous attractor-based accounts developed in spiking neural networks ^68,69^, while relying instead on the intrinsic dynamical richness generated by heterogeneous self-couplings and without assigning specialized computational roles to predefined neuronal subpopulations.

A distinctive feature of the hRNN is that contextual learning is achieved through relatively small modifications of the recurrent connectivity. Rather than constructing task representations from scratch, learning recruits and reshapes an already existing landscape of stable and metastable states. This parsimonious use of synaptic resources may provide two complementary advantages. First, previously acquired dynamical structures are less likely to be overwritten by new learning, thereby mitigating interference between consecutive tasks. Second, because task contexts can be encoded by small adjustments of an existing attractor scaffold, adaptation to novel situations may occur more rapidly than in architectures requiring extensive synaptic reorganization.

Interestingly, the behavior of the model closely resembles observations in both humans ^63^ and non-human primates ^64,70,71^, where rule switching emerges gradually following error feedback rather than through deterministic updating. In the hRNN, contextual changes are similarly mediated by transitions between attractor states, without requiring large-scale reorganization of synaptic connectivity. Moreover, contextual attractors are organized in a geometry of approximately parallel manifolds, reminiscent of the representational structures associated with abstraction and inferential reasoning in humans ^63,64^. We speculate that variability in the salience of error signals, together with other sources of neural stochasticity, may facilitate transitions between contexts by destabilizing the currently active attractor. The intrinsic multistability of hRNNs, characterized by a repertoire of attractors with heterogeneous degrees of stability, provides a natural substrate for expressing such variability. These fluctuations permit controlled exploration of alternative contextual states while preserving robust task representations, thereby exploiting the balance between flexibility and stability that characterizes the network dynamics. From this perspective, movement through the attractor landscape provides a mechanistic implementation of the exploration-exploitation trade-off underlying adaptive behavior ^72,73^.

Taken together, our results identify structural heterogeneity as a unifying mechanism linking scale-invariant memory and context-dependent computation. By generating both a broad spectrum of timescales and an intrinsic multistable landscape, heterogeneous recurrent networks support flexible behavior without requiring critical tuning. These findings raise the possibility that a common dynamical principle underlies the temporal richness, adaptive flexibility, and increasingly abstract representations observed across the cortical hierarchy.

## Methods

### Constructing heterogeneous RNNs with scale-invariant memory

We describe the procedure used to construct heterogeneous RNNs whose population activity decays as a power law after a polarized initial condition, in which all units are initialized at +1. This is achieved by first relating unit-specific self-couplings to local relaxation times and then choosing the distribution of self-couplings that yields the desired distribution of timescales.

#### Network dynamics: a rate model with heterogeneous self-couplings

We build upon the work in Ref. 45, which describes a neural network with *N* clusters of neuron populations, extending the previous seminal study in Ref. 53. In Ref. 45, intra-cluster connectivity is represented by self-connection variables *s*, introduced to capture strong local interactions that allow the network to exhibit bistability and chaotic dynamics. Extra-cluster connectivity, *W*_*ij*_ for *i* ≠ *j*, is modeled by Gaussian random connections. Similarly to the work in Ref. 47, as discussed in Sect. 2.1, we consider heterogeneous self-couplings *s*_*i*_ *≡ W*_*ii*_, *i* = 1, …, *N*. Differently from ^45,47^, we consider a rate model, describing firing rate variables instead of currents, to align more closely with a mean-field approach that describes leaky integrate-and-fire (LIF) neurons ^48–52^. These two models can be mathematically equivalent provided that the external input does not change in time ^74^. The network dynamics is then governed by the following differential equation:

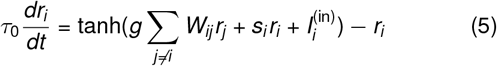

for *i* = 1, …, *N*, where *W*_*ij*_ s are drawn from a Gaussian distribution with mean 0 and variance 1*/N*, while *s*_*i*_ and *g* control the strengths of self- and inter-unit couplings, respectively. 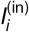 stands for the external input, and *τ*_0_ is the time constant representing the inertia of each neural unit. Unless otherwise stated, simulations used *N* = 500, recurrent gain *g* = 0.4, and single-unit time constant *τ*_0_ = 0.01 s.

#### Estimating local timescales from relaxation dynamics

To quantify how local timescales depend on the recurrent gain *g* and on the self-couplings *s*_*i*_, we analysed the relaxation dynamics of initially polarized networks. In this protocol, all units were initialized at *r*_*i*_ (0) = +1, and the subsequent dynamics were simulated in the absence of external input. For each fixed set of self-couplings 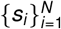, we averaged the activity over independent realizations of the off-diagonal connectivity matrix ***W***.

Averaging over connectivity realizations provides a direct estimate of how rapidly each unit forgets the polarized initial condition. For units in the bistable regime, activity switches between approximately symmetric states, so the averaged activity relaxes toward zero ^45^. For units in fluctuating or chaotic regimes, the activity also averages to zero because trajectories explore positive and negative firing-rate values over time. We therefore fitted the mean activity of each unit with an exponential relaxation,

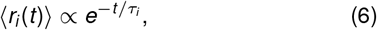

where *τ*_*i*_ is the characteristic relaxation time of unit *i*. This exponential approximation is equivalent to modeling the loss of memory of the initial condition as an effective Markovian relaxation process ^75^. Repeating this procedure across units with different self-couplings allowed us to obtain an empirical relationship *τ* (*s*) at fixed *g*. This relationship provides an operational link between the structural parameter *s* and the local timescale expressed by the network.

By linearity of the averaging operation, the population activity averaged over network realizations can be written as

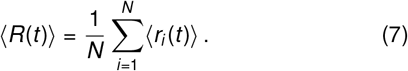

Thus, the population-level relaxation is a mixture of exponential decays with different characteristic timescales. Scale-invariant memory can therefore be obtained by choosing the distribution of self-couplings such that the induced distribution of relaxation times yields a power-law decay of ⟨ *R*(*t*) ⟩. In the following, we first derive the required distribution of timescales *P*_*τ*_ (*τ*), and then use the empirically fitted relationship *τ* (*s*) to determine the corresponding distribution of self-couplings *P*(*s*).

#### Deriving the target distribution of timescales

We next derive, in a similar spirit to Ref. 39, the distribution of relaxation times required to obtain a power-law decay of the population activity after a polarized initial condition. If the mean activity of each unit relaxes approximately exponentially, the population activity can be written as a mixture of exponential decays,

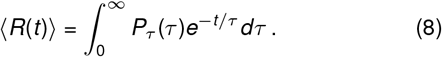

Introducing the change of variables *z* = 1*/τ*, this expression becomes

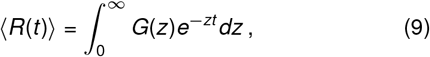

with

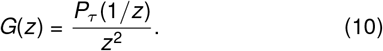

Thus, ⟨ *R*(*t*) ⟩ is the Laplace transform of *G*(*z*). To impose a target scale-invariant decay,

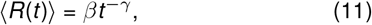

with *γ >* 0, we take the inverse Laplace transform and obtain

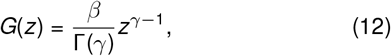

where Γ (*γ*) is the Gamma function ^76^. Transforming back to *τ* gives

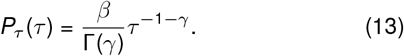

Therefore, an inverse-power-law distribution of relaxation times generates a power-law decay of the population activity.

In simulations, the range of available relaxation times is finite. We therefore use a truncated and normalized distribution on the interval [*τ*_min_, *τ*_max_],

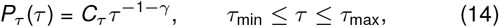

where

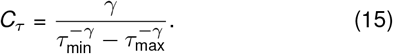

This finite support prevents the activity from diverging at *τ* = 0 and implies that the power-law decay is expected over the intermediate range *τ*_min_ ≪ *t* ≪ *τ*_max_.

Here, we use *γ* = 1 as a proof of concept. In this case,

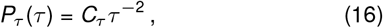

with

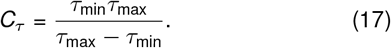

The resulting population activity is

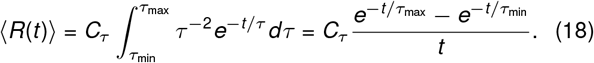

Thus, for *τ*_min_ ≪*t* ≪ *τ*_max_, the population activity follows

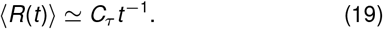

Other target exponents can be obtained by changing *γ*, including *γ* = 3*/*2, which is commonly associated with critical phenomena ^59^.

#### Deriving the distribution of self-couplings

By identifying the relationship between the relaxation time *τ*_*j*_ expressed by each unit *j* and its self-coupling *s*_*j*_, we can regulate the distribution of network timescales by adjusting the distribution of self-couplings, while keeping the recurrent gain *g* fixed. For our purposes, the precise functional form used to fit *τ* (*s*) is not essential. We fitted the empirical relationship using

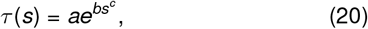

obtaining *a* = 0.0165 s, *b* = 1.02, and *c* = 2.99. The fit was performed in log-timescale space and provided an accurate description of the measured relationship over the parameter range considered (*R*^2^ = 0.98, Fig. 1e). Once *τ* (*s*) has been numerically established at fixed *g*, we can determine the distribution of self-couplings required to induce a target distribution of relaxation times, and therefore a power-law decay of the population activity (Fig. 1a).

Over the range of self-couplings used here, the fitted relationship *τ* (*s*) is monotonic, allowing us to obtain the distribution of self-couplings by a change of variables,

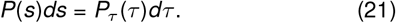

For the case *γ* = 1, derived above, we use

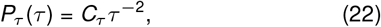

with

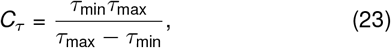

where *τ*_min_ = *τ* (*s*_min_) and *τ*_max_ = *τ* (*s*_max_). Since

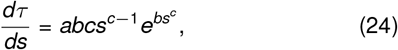

the induced distribution of self-couplings is

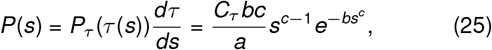

for *s*_min_ *≤ s ≤ s*_max_.

To sample self-coupling values from this distribution, we used a standard Monte Carlo sampling from the cumulative distribution

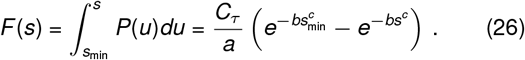

Taking *Y* as uniformly distributed in [0, 1], the corresponding self-coupling is obtained as

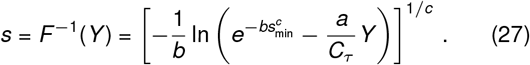

For the simulations shown in Fig. 1, we used *s*_min_ = 0.7 and *s*_max_ = 1.7. The resulting distribution *P*(*s*) is not intended as a mechanistic law for self-couplings. Rather, it is the distribution required, given the fitted relationship *τ* (*s*), to induce the target distribution of relaxation times and thereby produce a scale-invariant decay of the population activity. A mechanistic interpretation of the fitted *τ* (*s*) relationship is developed in the following section.

#### Simulation details for Fig. 1

For the simulations shown in Fig. 1, networks contained *N* = 500 units with recurrent gain *g* = 0.4 and single-unit time constant *τ*_0_ = 0.01 s. Self-couplings were sampled in the range *s*_min_ = 0.7 to *s*_max_ = 1.7 using the distribution derived above. All units were initialized at *r*_*i*_ (0) = +1, and the dynamics were simulated in the absence of external input. Dynamics were integrated for 2 s using an explicit Euler scheme with time step *dt* = 10^−3^ s. Mean activities were computed by averaging over 500 independent realizations of the synaptic matrix ***W*** (with null diagonal elements), keeping the self-couplings fixed. Relaxation times *τ*_*i*_ were obtained by fitting the connectivity-averaged activity of each unit with an exponential decay over the full simulated interval.

#### Asymptotic stability and robustness

To characterize the asymptotic dynamics of the heterogeneous network, we simulated 100 independent realizations of ***W*** (with null diagonal elements), keeping the self-couplings fixed, from the polarized initial condition *r*_*i*_ (0) = +1 for up to 10^4^ s. Convergence to a fixed point was determined from the residual (Fig. S1b)

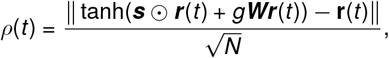

which was required to remain below 10^*−*7^ for at least 20 s. The convergence time was defined as the first crossing of this threshold. We estimated the two largest Lyapunov exponents (Fig. S1c) over a 300 s window using the standard algorithm in Ref. 77. To assess robustness, we repeated the analysis for *g* ∈ [0.38, 0.42], averaging over independent connectivity realizations while keeping the self-coupling values fixed.

### Effective-noise approximation of recurrent input fluctuations

To test whether the recurrent interactions of the heterogeneous network can be approximated by an effective stochastic drive, we compared the coupled network used in Fig. 1 with a fully decoupled network driven by independent Ornstein– Uhlenbeck (OU) processes (Fig. 2a). Both networks contained *N* = 500 units, used the same self-couplings {*s*_*i*_*}* distributed as in Fig. 1, and had intrinsic time constant *τ*_0_ = 0.01 s. In the coupled network, the recurrent gain was fixed to *g* = 0.4. All simulations were initialized from the polarized state *r*_*i*_ (0) = +1 for all units and integrated for 2 s with time step *dt* = 10^−3^ s. Averages were computed over 500 independent realizations of the off-diagonal recurrent connectivity in the coupled network and, in the decoupled network, over independent realizations of the OU process provided as inputs.

#### Coupled network and recurrent currents

In the coupled configuration, the dynamics are

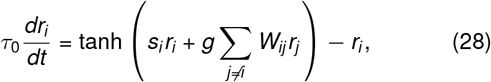

or in matrix form

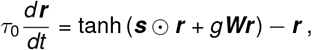

where the off-diagonal recurrent weights *W*_*ij*_, with *i* ≠ *j*, are drawn independently from *N*(0, 1*/N*) and with null diagonal elements (*W*_*ii*_ = 0). For each realization, we recorded the recurrent input current received by each unit, excluding the self-coupling contribution, 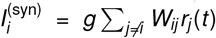 with a standard deviation 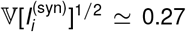. This value was not imposed in the OU-driven network, but was used as a consistency check for the fitted OU amplitude.

#### OU-driven decoupled network

In the decoupled configuration, all off-diagonal recurrent interactions were removed while preserving the same self-couplings *{s*_*i*_}:

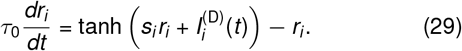

The recurrent input to each unit was replaced directly by an independent OU process,

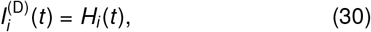

with

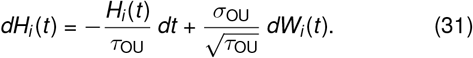

Here, *W*_*i*_ (*t*) is an independent standard Wiener process for unit *i*, so that ⟨*dW*_*i*_ (*t*) ⟩ = 0 and ⟨*dW*_*i*_ (*t*)*dW*_*j*_ (*t*) ⟩= *δ*_*ij*_ *dt*. With this convention, each OU process has stationary variance

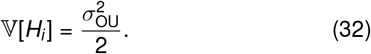

In simulation, the OU processes were initialized from their stationary distribution,

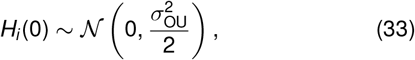

and simulated with the Euler–Maruyama method.

#### Fitting the OU amplitude

We fixed the OU correlation time to the intrinsic relaxation time of the rate units,

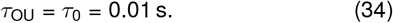

This choice follows the colored-noise theory of bistable neuronal populations, for which an exact reduction is available when the correlation time of the input noise matches the intrinsic timescale of the rate unit ^60^. We then fitted only the OU amplitude *σ*_OU_.

The fitted value was obtained by a one-dimensional grid search over *σ*_OU_, selecting the value that minimized the mean squared difference between the mean population activities of the coupled and decoupled networks, ⟨*R*_*C*_⟩ and ⟨*R*_*D*_⟩ respectively, over the relaxation window *T*_fit_ = [0.003, 1.9] s,

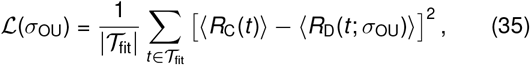

where

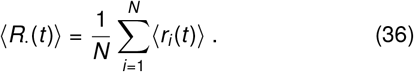

The fitted OU amplitude was

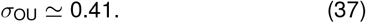

With our convention for the OU process, this corresponds to a stationary current standard deviation

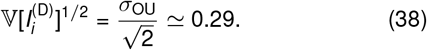

This value is close to the measured standard deviation of the recurrent currents in the coupled network, *≃* 0.27, supporting the interpretation that the fitted OU drive provides input fluctuations of comparable magnitude to those generated internally by the recurrent network.

#### Theoretical prediction for bistable units

To compare the measured relaxation times with the theory of bistable rate units driven by colored synaptic noise, we used the analytical prediction derived in Ref. 60. This comparison was restricted to units in the bistable regime (*s*_*i*_ *>* 1). The theoretical prediction applies to a single rate unit driven by a colored input current whose correlation time matches the single-unit time constant. We therefore used the case *τ*_OU_ = *τ*_0_, consistent with the OU-driven decoupled network described above.

For each value of *s >* 1, we evaluated the theoretical timescale *τ*_th_(*s*) using the expression derived in ^60^. The noise-amplitude parameter entering this expression was set directly to the amplitude of the OU current driving each unit, i.e., *σ*_OU_ *≃* 0.41. The resulting theoretical prediction was compared with the relaxation times extracted from the decay of the mean activity in Fig. 2d.

### RNNs performing short-term memory task

Performance comparisons on the memory task were conducted on the hRNN, the homogeneous RNN and the ESN. All networks comprised *N* = 500 units; the hRNN and the homogeneous RNN also shared the same recurrent gain *g* = 0.4 and time constant *τ*_0_ = 0.01 s. The hRNN self-couplings *s*_*i*_ were distributed as described in the previous sections, between *s*_min_ = 0.7 and *s*_max_ = 1.7. For a fair comparison, the homogeneous self-coupling was set to the mean of the Hrnn distribution, 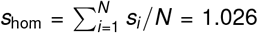. For the classical ESN we used *g*_ESN_ = 0.9999, i.e., a spectral radius at the edge of stability, *s*_*i*_ = *s*_ESN_ = 0 and *τ*_0_ = 0.04 s.

We presented each network with a sequence of six successive, mutually orthogonal input patterns targeting six disjoint subsets that partitioned the network, each containing approximately one sixth of the total units. Each stimulus consisted of a strong positive external input applied for 10 ms, saturating the transfer function of the targeted units at +1 and thereby driving their activity toward this value, without clamping it. Units outside the stimulated subset received no external input and continued to evolve according to the network dynamics. Successive stimuli were separated by an inter-stimulus delay *T*_sep_, measured from stimulus offset to the next onset. We tested *T*_sep_ = 0.01, 0.05, and 0.2 s. For each ***W*** realization, five initial stimulation cycles were discarded as a transient, followed by ten consecutive recorded cycles, with no reset of the network state or additional pause between cycles. For each stimulus, we defined the evoked population activity vector as the time average of the network activity **r**(*t*) ∈ ℝ^*N*^ over the inter-stimulus delay following that stimulus. For each *T*_sep_, we then computed the Pearson correlation coefficient across units between the vectors evoked by pairs of stimuli, as a function of their ordinal distance in the sequence. At each stimulus distance *d* in the sequence, correlations were averaged over all pairs of stimuli within the recorded cycles and then over 50 realizations of the synaptic matrix ***W***, keeping the self-couplings fixed. The ESN time constant was chosen so that its correlations at *T*_sep_ = 0.01 s closely matched those of the homogeneous network (top row of Fig. 3c).

Since *ρ* is symmetric in its two arguments, correlations at signed distances *±d* coincide; both signs are shown in Fig. 3b for visual clarity.

### Reservoir computing approach

Reservoir computing is a powerful computational framework that utilizes RNNs to perform tasks and emulate dynamical systems^44,62,78^. It is characterized by fixed, random internal connections. The network transforms input data through its complex dynamics, and the resulting “reservoir” of internal states is fed into a simpler readout layer to generate the final output. The most widely known form of reservoir computing is the echo-state network (ESN) ^62^, although numerous variants have been proposed over the years ^79–82^. ESNs are RNNs whose units are designed to represent small, homogeneous clusters of neurons, akin to those observed in cortical circuits ^48–52^. Each of the *N* units, represented as *r*_*i*_ (*t*), follows the firing rate equation

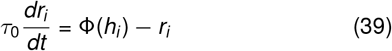

Here, *r*_*i*_ (*t*) is the activity of the *i*-th unit, *τ*_0_ is a time constant, and Φ (*h*_*i*_) = tanh(*h*_*i*_) is a sigmoidal function that governs how each unit responds to its inputs. The input to each unit, *h*_*i*_ (*t*), is a weighted combination of signals from other units within the network and any external input it receives. The internal input comes from the synaptic matrix ***W***, which delineates the connections between units and defines the specific reservoir model. In ESNs, this matrix is randomly generated from a Gaussian distribution. In our exploration, we examined four scenarios: null, constant, or heterogeneous self-interactions (the diagonal of the matrix), along with a case involving heterogeneous *τ*_0_ values.

The external input is represented by 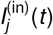 and is linked to some observable data fed into the system, denoted as ***u***(*t*) ∈ ℝ ^*P*^. This input data is connected to the network through random weights, so that 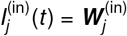, where 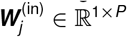 is he *j*-th row of the input matrix ***W*** ^(in)^ *∈* ℝ ^*N × P*^.

In the reservoir computing framework, during training, the network collects the time series data of each unit into a matrix ***R*** ∈ ℝ^*N × T*^, where each row corresponds to the activity of a unit over time. Similarly, the target outputs are represented in a matrix ***Y*** ∈ ℝ ^*M × T*^, with each row representing a target output for the system to learn. Learning consists in finding the readout weights ***W*** ^(out)^ ∈ ℝ ^*M × N*^ optimally mapping the internal states and the outputs. They are the result of the ridge regression

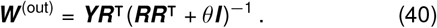

In this equation, *θ* is a regularization parameter to prevent overfitting, and ***I*** is the identity matrix. When the target output coincides with the external input ***u***(*t*), the trained readout layer can be used to predict future inputs based on the current state of the RNN. This procedure effectively corresponds to updating the synaptic matrix with a correction term given by Δ***W*** = ***W*** ^(in)^***W*** ^(out)^, providing a straightforward rule for adjusting synaptic strengths.

### Heterogeneous RNNs solving WCST task

In Sect. 2.4, we examine heterogeneous neural networks in reservoir computing tasked with performing the Wisconsin Card Sorting Test (WCST). Given a central card and a pair of oriented cards, the system must respond based on the current context, which evolves over time with a certain probability.

We consider two contexts, each containing two items. This can be conceptualized as four possible cards, each represented by two attributes: color and shape. For instance, the cards could be yellow-triangle, yellow-circle, green-triangle, and green-circle, as illustrated in Fig. 4a. The governing rule can either be “color” or “shape”. If the context is based on color and the central card is yellow, the correct response will be to select the yellow card from the pair of oriented cards.

Each item corresponds to a stimulus with varying weights on the network units. All external stimuli provided to the network are in the form of smoothed square waves. The pair of items is differentiated as one positive and one negative, with the response determined by the sign of the chosen card.

A trial begins with the stimulation of the central card, which lasts for 0.09 s. After a 0.01 s delay, the currents for the pair of cards are introduced, also lasting 0.09 s. After 0.03 s during which the oriented cards are visible, the network must generate a response. If the response is correct, the network is allowed to relax for the next 0.1 s. However, if the response is incorrect, a positive error signal is sent with a 0.01 s delay, lasting 0.06 s, to notify the system.

During the learning phase, an additional context current (positive or negative, depending on the prevailing rule) is applied. A linear readout is trained to reproduce the optimal response: the correct response, except on the trial of a rule switch, where it is the correct response of the previous rule, since the switch cannot be anticipated. A second readout is trained to autonomously generate the context signal, which is subsequently fed back into the network as feedback.

For the WCST task, we conduct 20 transient trials for dynamic stabilization, which are excluded from the readout learning process. These are followed by 80 trials for learning and 100 trials for testing. The rule may change between trials with a probability of 20%. Notably, an incorrect response during a trial where a rule change occurs is inevitable, and is therefore not penalized in accuracy calculations.

We implemented two types of heterogeneity in networks comprising 500 units, with internal connectivity strength set to *g* = 0.4, and external input strengths of *g*_error_ = 1.75 and *g*_card_ = 1.0. The networks differ in terms of self-coupling and time constants. The hRNNs, identical to those used in Sect. 2.3, feature self-coupling values ranging from *s*_*i*_ *∈* [0.7, 1.7], whereas the *τ*_0_-networks have no self-coupling (*s*_*i*_ = 0). Time constants are uniform across all units in the hRNNs (*τ*_0_ = 0.01 s). In contrast, the *τ*_0_-networks have time constants uniformly distributed within *τ*_0_ ∈ [0.01, 0.05] s as in Ref. 81. The network dynamics are integrated using the first-order Euler method with a time step of *dt* = 0.001 s.

Finally, the readout layer is trained using ridge regression, with a regularization coefficient of *θ* = 0.01.

#### Autocorrelation of the closed-loop network

After training, the closed-loop network, with no external input, was started from 100 random initial states (Gaussian, rescaled to max_*i*_ |*r*_*i*_ (0) | = 1) and simulated for 10 s. For each start we computed *C*(*t*) = **r**(0) *·* **r**(*t*)*/*( ‖**r**(0) ‖ ‖ **r**(*t*) ‖) every 5 ms and averaged it over the starts (Fig. 5e, mean *±* SEM over the 50 networks). A power law *C*(*t*) = *αt*^*−γ*^ was fitted by least squares from *t* = 0.02 s on, excluding the initial plateau.

#### Accuracy around a rule switch

To compare the behavior of the network with that of humans (Fig. 4f), we trained 200 hRNNs with *g*_fb_ = 0.8 as described above, with the full error signal, and tested each of them four times, halving each error signal of the test with probability *p* = 0 or 0.75 (one independent draw per trial). The connectivity, the task sequence and the training of each network were the same for every *p*, so that the four conditions differ only in the strength of the error signals. In this analysis, accuracy is the fraction of correct responses under the current rule, i.e., of responses not followed by an error signal. We considered rule switches occurring between two blocks of at least four test trials (977 switches across 200 networks, identical for all values of *p*). For each network, we computed the accuracy on the last trial before each switch and on the first four trials after it, averaging over all switches. We report the mean *±* SEM of these values across networks.

## Code availability

All code required to reproduce the results presented in this study is publicly available at https://github.com/gdianto/ScaleInvariantRNN.

## Acknowledgments

This work was partially supported by the EU Horizon Europe programme under the Marie Skłodowska-Curie grant agreement No 101126667 for the Ramon Llull-AIRA Postdoctoral Programme to GDA, and by the Italian National Recovery and Resilience Plan (PNRR), M4C2, NextGenerationEU (Project IR0000011, CUP B51E22000150006, “EBRAINS-Italy”) to MM.

