## Supplemental Figure S1 for "Heterogeneous self-couplings enable scale-invariant memory and cognitive flexibility without criticality"

### Supplementary materials

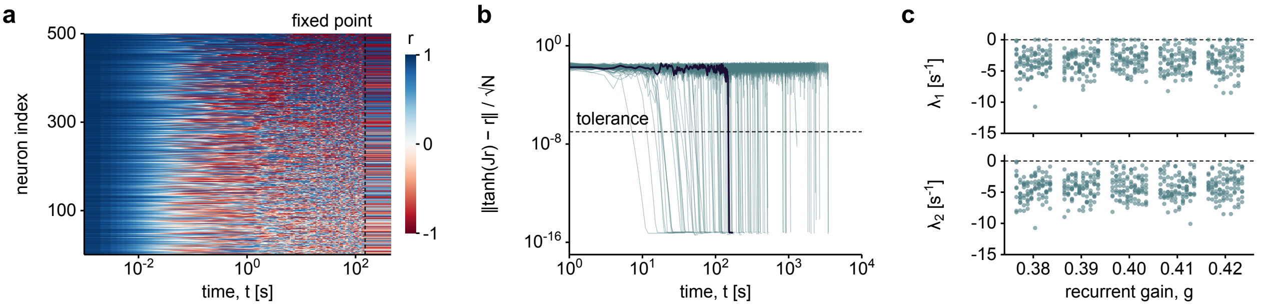


Figure S1: **Asymptotic stability and robustness of the heterogeneous RNN. a,** Representative network activity starting from the polarized initial condition $r_i(0) = 1$, showing irregular transient dynamics followed by convergence to a fixed point. Time is shown on a logarithmic scale. **b,** Evolution of the fixed-point residual $\rho(t)$ across independent connectivity realizations that converged to stable fixed points. The black line corresponds to the network realization shown in panel a. The dashed horizontal line indicates the convergence threshold $\rho = 10^{-7}$. A network is considered converged when the residual remains below this threshold for at least 20 s. Here, $J$ denotes the full connectivity matrix, with $J_{ii} = s_i$ and $J_{ij} = gW_{ij}$ for $i \neq j$. **c,** First and second Lyapunov exponents ($\lambda_1$ and $\lambda_2$) estimated across network realizations at recurrent gains $g = 0.38$–$0.42$, corresponding to $\pm 5\%$ variations around the reference value $g = 0.4$. Each point at fixed $g$ represents an independent network realization. Dashed horizontal lines indicate zero. Network parameters are as in Fig. 1.
